# A new high-quality genome assembly and annotation for the threatened Florida Scrub-Jay (*Aphelocoma coerulescens*)

**DOI:** 10.1101/2024.04.05.588142

**Authors:** Faye G. Romero, Felix E.G. Beaudry, Eyvind Hovmand Warner, Tram N. Nguyen, John W. Fitzpatrick, Nancy Chen

**Affiliations:** Department of Biology, University of Rochester. Rochester, NY, USA; Ontario Institute for Cancer Research. Toronto, ON, Canada; Department of Ecology & Evolutionary Biology, Cornell University. Ithaca, NY, USA; Cornell Lab of Ornithology, Cornell University. Ithaca, NY, USA

**Keywords:** Florida Scrub-Jay, Corvidae, de novo genome assembly, linkage map, pedigree

## Abstract

The Florida Scrub-Jay (*Aphelocoma coerulescens*), a Federally Threatened, cooperatively-breeding bird, is an emerging model system in evolutionary biology and ecology. Extensive individual-based monitoring and genetic sampling for decades has yielded a wealth of data, allowing for the detailed study of social behavior, demography, and population genetics of this natural population. Here, we report a linkage map and a chromosome-level genome assembly and annotation for a female Florida Scrub-Jay made with long-read sequencing technology, chromatin conformation data, and the linkage map. We constructed a linkage map comprising 4,468 SNPs that had 34 linkage groups and a total sex-averaged autosomal genetic map length of 2446.78 cM. The new genome assembly is 1.32 Gb in length, consisting of 33 complete or near-complete autosomes and the sex chromosomes (ZW). This highly contiguous assembly has an N50 of 68 Mb and a Benchmarking Universal Single-Copy Orthologs (BUSCO) completeness score of 97.1% with respect to the *Aves* database. The annotated gene set has a BUSCO transcriptome completeness score of 95.5% and 18,051 identified protein-coding genes, 92.2% of which have associated functional annotations. This new, high-quality genome assembly and linkage map of the Florida Scrub-Jay provides valuable tools for future research into the evolutionary dynamics of small, natural populations of conservation concern.

**ARTICLE SUMMARY:** We present a new high-quality genome assembly and annotation for the Florida Scrub-Jay (*Aphelocoma coerulescens*), a Federally Threatened bird species. In comparison to other genome assemblies of this species, our assembly is the first to be made using long-read sequencing technology and is the first generated from a female individual. We also constructed the first linkage map for this species using a population pedigree. Our genome assembly is highly contiguous and is of similar quality to other bird genome assemblies.

## INTRODUCTION

The Florida Scrub-Jay (*Aphelocoma coerulescens*) is a Federally Threatened, cooperatively-breeding bird endemic to the U.S. state of Florida (Figure 1A) (Woolfenden and Fitzpatrick 1984). This species has been in decline due to anthropogenic development and fire suppression, and currently exists in small, locally isolated populations across the state (Boughton and Bowman 2011). The Florida Scrub-Jay is intensively monitored throughout its range with several well-characterized natural populations, including a long-term study at Archbold Biological Station in Venus, FL. All individuals in this population have been uniquely banded and monitored since 1969, resulting in a 16-generation pedigree with near-complete fitness data (Figure 1B) (Woolfenden and Fitzpatrick 1984). This robust dataset has led to foundational knowledge in the behavior, demography, and life history of cooperative breeders (Woolfenden and Fitzpatrick 1984), with further work shedding light on social and environmental effects on lifetime fitness (Mumme *et al*. 2015) and the causes and consequences of dispersal and immigration (Coulon *et al*. 2010; Aguillon *et al*. 2017; Suh *et al*. 2020, 2022; Summers *et al*. 2024). Studies of other populations of Florida Scrub-Jays have contributed to our understanding of the negative impacts of suburbanization (Thorington and Bowman 2003; Coulon *et al*. 2012), population dynamics (Breininger *et al*. 1999; Breininger and Carter 2003), and the repercussions of translocations (Linderoth *et al*. 2023). Exhaustive genetic sampling of thousands of individuals at Archbold Biological Station has also allowed for the rare opportunity to study the evolution of small, natural populations, such as genetic population structure (Coulon *et al*. 2008, 2010), allele frequency changes (Chen *et al*. 2019), and the genetic consequences of inbreeding and reduced immigration (Chen *et al*. 2016; Nguyen *et al*. 2022), all of which are imperative to understand in the face of habitat fragmentation and the loss of genetic diversity in species worldwide (Thomas *et al*. 2004). To better characterize the genetic variation present in the Florida Scrub-Jay, we must have a high-quality reference genome as a point of comparison.

**Figure 1.**
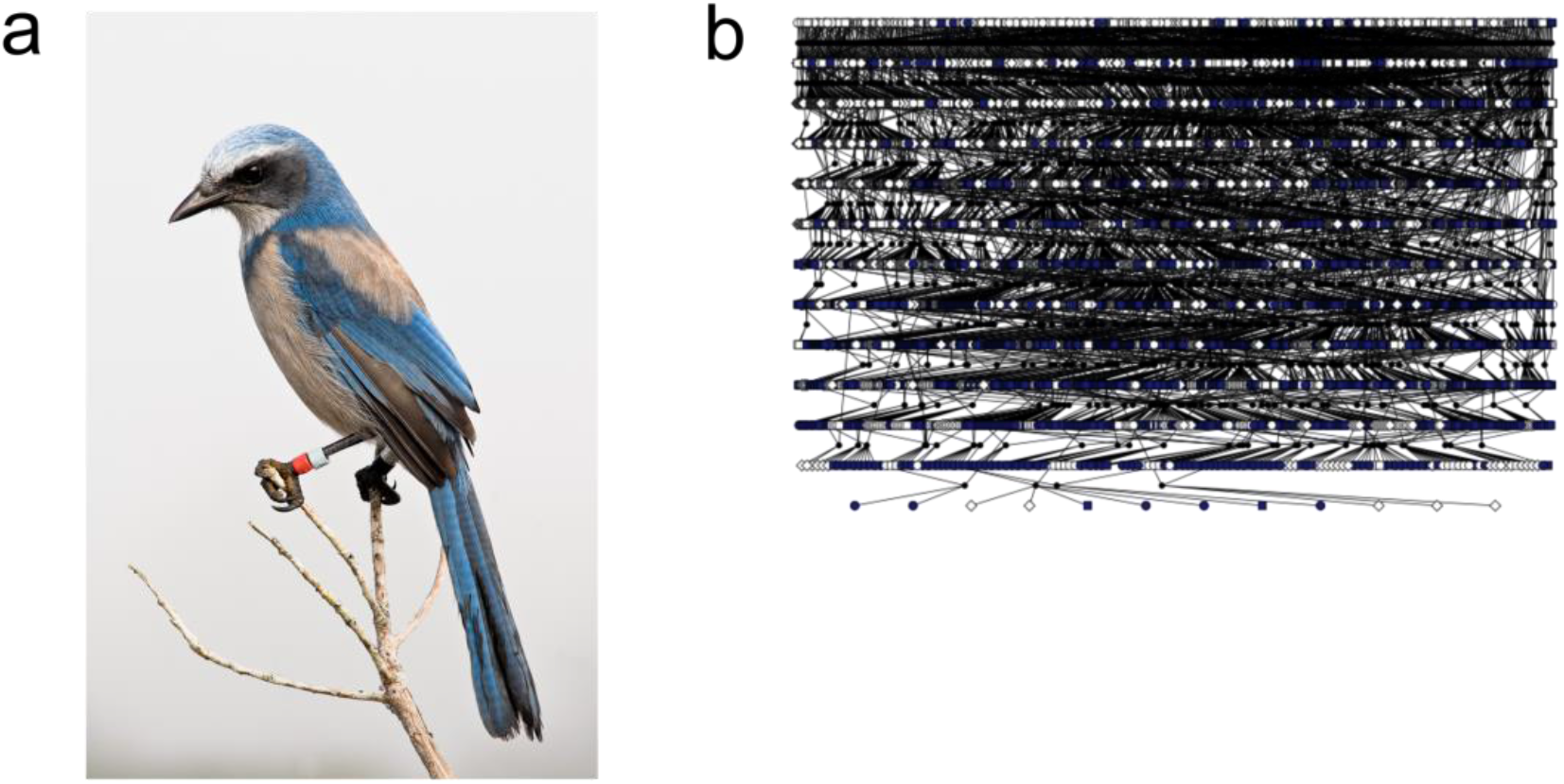
Image and population pedigree of the Florida Scrub-Jay (*Aphelocoma coerulescens*). (A) A banded Florida Scrub-Jay from the long-term demographic study at Archbold Biological Station. Photo courtesy of Reed Bowman. (B) The population pedigree for Florida Scrub-Jays at Archbold Biological Station from 1969-2013. Blue symbols indicate individuals who have been genotyped. N ∼ 14,000.

A reference genome for a male Florida Scrub-Jay was published as part of the Bird 10,000 Genomes Project (version 1; v1) (Feng *et al*. 2020) and was further improved and scaffolded with the aid of Hi-C reads (version 2; v2) (Driscoll & Beaudry, *et al*. 2021). However, these assemblies were generated with Illumina short-read data, which may not have captured the full scope of genomic information, such as highly repetitive regions (Treangen and Salzberg 2012). Additionally, as birds have a ZW sex-determination system in which females are the heterogametic sex, these assemblies are missing the W chromosome. Here, we present a new chromosome-level genome assembly for a female Florida Scrub-Jay generated with long-read sequencing technology, chromosome conformation data, and a linkage map. The version 3 (v3) assembly, which is 1.32 Gb long, adds 260 Mb in length to the previous reference genomes and contains 4 newly identified chromosomes to the Florida Scrub-Jay, including the W chromosome. We also provide annotations of repetitive and gene content, evaluations of the quality and contiguity of the data presented, and the first linkage map for this species.

## MATERIALS AND METHODS

### Sampling and genome sequencing

We collected fresh blood via venipuncture from an inbred, adult female Florida Scrub-Jay at Archbold Biological Station, Venus, FL (approved by Cornell University Institutional Animal Care and Use Committee (IACUC 2010-0015), authorized by permit no. TE824723-8 issued by the US Fish and Wildlife Service, banding permit no. 07732 issued by the US Geological Survey, and permit no. LSSC-10-00205 issued by the Florida Fish and Wildlife Commission). The University of Delaware DNA Sequencing & Genotyping Center extracted DNA from the blood sample using a High Molecular Weight extraction protocol, then prepared a Pacific Biosciences (PacBio) library and sequenced it on 3 SMRT Cells (Sequel IIe system).

We also performed 20x coverage whole-genome resequencing for 25 males and 25 female Florida Scrub-Jays, including the parents of the individual we sampled for PacBio sequencing. We extracted DNA from archived blood samples stored in Queen’s lysis buffer using Qiagen DNeasy Blood and Tissue kits and sent DNA to Novogene (Sacramento, CA, USA) for PCR-free library preparation and 150 bp paired-end sequencing on an Illumina NovaSeq6000 platform.

### Linkage map generation

We created a linkage map using CRI-MAP v. 2.507 (Green *et al*. 1990) and the CRIGEN package (Liu and Grosz 2006). We used data from a previous study that genotyped 3,838 individuals at 12,210 SNPs using a custom Illumina iSelect Beadchip (Chen *et al*. 2016). We trimmed our pedigree to include only completely genotyped trios, then split it into 32 three-generation sub-families of ∼100 individuals each using the *crigen* function. Using a subset of 3,424 informative SNPs (1 SNP per scaffold of Florida Scrub-Jay genome v1; Feng *et al*. 2020), we created a sparse (pre-framework) map. We calculated pairwise LOD scores using *twopoint* and assigned markers to linkage groups using the *autogroup* function. *Autogroup* uses an iterative process to assign markers to linkage groups with four levels of increasing stringency.

The parameters we used for minimum LOD score, minimum number of informative meioses, maximum number of shared linkages, and minimum linkage ratio were: level 1 (100, 2.0, 2, 0.9), level 2 (50, 1.5, 3, 0.7), level 3 (10, 1.0, 5, 0.6), and level 4 (5, 0.4, 6, 0.5). We labelled linkage groups based on alignments with the Zebra finch genome (NCBI accession: GCA_000151805.2). Then, to construct each linkage group, we identified haplogroups using the *hap* function and ran *build* four times with different starting markers and a threshold of LOD > 5. We chose the longest map as the pre-framework map for each linkage group. To check marker order, we permuted up to five adjacent markers with the function *flips* to look for alternative marker orders with higher likelihood and iteratively updated the marker order until no better orders were found.

To expand the pre-framework map, we ran *twopoint* and *autogroup* on the full SNP set as above to assign the remaining markers to linkage groups. For each linkage group, we added markers onto the pre-framework map using *build* with a threshold of LOD > 5 and confirmed marker order with *flips*. For three linkage groups (LG 34, 36, and chr Z), we added additional markers using *build* with a threshold of LOD > 3 and checked marker order with *flips*. When linkage groups had multiple equivalent best orders, we either removed markers with multiple potential orders or picked the order that was most consistent with the physical map. Finally, we used *fixed* to output the maximum likelihood recombination fractions and map distances for the sex-averaged map (setting SEX_EQ to 1) and the sex-specific map (setting SEX_EQ to 0). We used crimaptools v0.1 (https://github.com/susjoh/crimaptools, (Johnston *et al*. 2016)) to parse output files.

### *De novo* genome assembly

To assemble the genome, we first used Cutadapt v. 2.3 (Martin 2011) to identify and discard PacBio HiFi raw reads with adapter sequences. Next, we created primary and alternate draft assemblies using hifiasm v. 0.16.1 (Cheng *et al*. 2021) in HiFi-only mode with default parameters. We also ran Hifiasm in trio-binning mode, which leverages short-read data from the reference individual’s parents to generate haplotype-resolved assemblies. We prepared the maternal and paternal reads by trimming adapters and filtering for quality with fastp v. 0.21.0 (Chen *et al*. 2018), merging paired-end reads with PEAR v. 0.9.11 (Zhang *et al*. 2014a), and building a k-mer hash table for each set of reads using yak v. 0.1 (Li 2020). We compared the quality and contiguity of the four draft assemblies (primary, alternate, maternally-resolved haplotype, paternally-resolved haplotype) with Quast v. 5.0.2 (Mikheenko *et al*. 2018) and BUSCO v. 5.2.2 (using the aves_odb10 and eukaryote_odb10 databases; Manni *et al*. 2021) and moved forward with the most contiguous assembly (the primary assembly). To further scaffold the genome, we used the 95.7 Gb of Hi-C reads generated by Dovetail Genomics for the v2 Florida Scrub-Jay genome assembly (Driscoll & Beaudry, *et al*. 2021). We used the Arima Hi-C mapping pipeline (https://github.com/ArimaGenomics/mapping_pipeline) to map the paired-end Hi-C reads to the hifiasm primary assembly and SALSA v. 2.3 (Ghurye *et al*. 2017) to join the contigs into scaffolds. We visualized the Hi-C contact map and manually curated the scaffolds to generate a chromosome-level genome assembly using the *juicer.sh*, *run-assembly-visualizer.sh*, and *run-asm-pipeline-post-review.sh* scripts from the Juicer v. 1.6 pipeline (Durand *et al*. 2016). To further improve contiguity, we ordered and oriented scaffolds given positional evidence from our linkage map using ALLMAPS v. 1.3.7 from the JCVI Utilities Library (Tang *et al*. 2015). If any marker locations in the linkage map conflicted with the ordering of contigs joined during the SALSA or Juicer scaffolding steps, we manually broke those scaffolds in disagreement and iteratively ran ALLMAPS until the genetic and physical positions for each linkage group were in concordance. We next screened the genome for organismal contaminants using the BlobToolKit suite v. 3.1.0 (Challis *et al*. 2020) with the *--busco*, *--hits*, and *--cov* flags, and conducted a BLAST search (BLAST v. 2.10.0+; Camacho *et al*. 2009) against the publicly available Florida Scrub-Jay mitochondrial genome sequence (NCBI accession NC_051467.1) to identify and remove any mitochondrial contaminants in the assembly. Finally, we numbered the linkage groups according to homology with the Zebra finch reference genome (*Taeniopygia guttata*; bTaeGut1.4.pri, NCBI accession GCA_003957565.4). Full descriptions of all software and options used for *de novo* genome assembly are available in Table S1.

### Sex chromosome identification

We identified Z- and W-linked scaffolds using a two-pronged approach: relative read depth and sequence homology to other bird species. Using whole-genome sequence data from 50 Florida Scrub-Jays, 25 male and 25 female, we followed the basic methodology from the findZX pipeline (Sigeman *et al*. 2022), a computational pipeline for sex chromosome identification. We processed each set of raw reads as follows: 1) trimmed adapter sequences and low-quality reads using Trim Galore (Krueger *et al*. 2023), 2) mapped the raw reads to the genome assembly using BWA-MEM (Li 2013), 3) filtered for read pairs that completely mapped in the expected orientation with a mapping quality greater than 20 using samtools *view* (Danecek *et al*. 2021), 4) marked and removed duplicate reads using sambamba (Tarasov *et al*. 2015), 5) filtered for reads with an edit distance of less than or equal to 2 using bamtools *filter* (Barnett *et al*. 2011), and 6) calculated per-basepair read depth and average read depth per scaffold across the genome, ignoring repetitive and low complexity regions, using a custom Bash script and samtools *mpileup* (Danecek *et al*. 2021). We then compared the average read depth of each scaffold between males and females in R using a series of t-tests with significance values Bonferroni-corrected for multiple comparisons. We putatively assigned scaffolds with significantly different average read depths as Z-linked if read depth was higher in males than in females and W-linked if read depth was higher in females than in males. We confirmed sex chromosome assignments by aligning each putatively sex-linked scaffold to its appropriate sex chromosome in the Zebra finch reference genome and checking for sequence homology. As an additional check, we aligned W-linked scaffolds to the paternally-resolved haplotype assembly to confirm that they are missing from the male (ZZ) assembly. Finally, we labeled scaffolds as Z-linked or W-linked if they yielded both a significant t-test and displayed sequence homology to the Zebra finch sex chromosomes. Full software parameters are available in Table S1.

### Genome annotation

#### Repetitive element annotation

To annotate repetitive content across the genome, we first constructed a custom repeat library for the Florida Scrub-Jay using RepeatModeler v. 2.0.4 (Flynn *et al*. 2020) with the *-LTRStruct* flag. We then merged this library with a curated avian repeat library (Peona *et al*. 2021b) and a curated repeat library of the closely-related Steller’s Jay (*Cyanocitta stelleri*; Benham *et al*. 2023). We used RepeatMasker v. 4.1.4 (Smit *et al*. 2013) to identify repetitive regions across the genome with the *-s* and *-xsmall* flags to implement a slow search and softmask the genome, respectively. To assess how sequencing technologies (*i.e.*, short-read versus long-read) impacted repeat annotation, we compared the total counts and median lengths of transposable element (TE) superfamilies (Kapitonov and Jurka 2008) across the v2 and v3 assemblies using Wilcoxon rank sum tests.

#### Gene prediction and functional annotation

Robust and highly confident gene prediction leverages both RNA-seq and protein data. As such, we used the BRAKER pipeline v. 3.0.6, which integrates both types of data to train and execute the GeneMark-ETP and AUGUSTUS gene prediction tools (Stanke *et al*. 2006, 2008; Gotoh 2008; Iwata and Gotoh 2012; Buchfink *et al*. 2015; Hoff *et al*. 2016, 2019; Kovaka *et al*. 2019; Pertea and Pertea 2020; Brůna *et al*. 2021; Bruna *et al*. 2024). We obtained trimmed and filtered 2×101bp RNA-seq reads from liver, heart, and kidney samples from one male and one female, as well as ovary samples from the female (Driscoll & Beaudry, *et al*. 2021). Next, we used STAR v. 2.7.3 (Dobin *et al*. 2013) to align the RNA-seq reads to the softmasked genome assembly and Picard v. 2.27.4 (Broad Institute 2019) to assign read groups to each sample. We then executed BRAKER with the softmasked genome, aligned RNA-seq reads, and the *Vertebrata* protein sequence database from OrthoDB v.11 (Kuznetsov *et al*. 2023) as input data. We used InterProScan v. 5.65-97.0 (Jones *et al*. 2014) and a protein BLAST search against the Swiss-Prot database (The UniProt Consortium 2019) to assign functional annotations to the resulting gene set. Finally, we combined the outputs of BRAKER and InterProScan into a consensus gene annotation using the AGAT v. 1.2.0 suite of tools (Dainat 2023). Full descriptions of all software and options used for genome annotation are available in Table S1.

### Genome completeness assessment

We used QUAST to calculate basic assembly quality statistics and BUSCO to assess expected gene content and completeness across the genome (as above). To explore synteny across the *Aves* group, we used minimap2 v. 2.26 (Li 2018) to generate whole genome-whole genome alignments of our assembly with publicly available Zebra finch, chicken (*Gallus gallus*; GGswu, NCBI accession GCA_024206055.2), New Caledonian crow (*Corvus moneduloides*; bCorMon1.pri, NCBI accession GCA_009650955.1), and the closely related California Scrub-Jay (*Aphelocoma californica*; bAphCal1.0.hap1, NCBI accession GCA_028536675.1) genomes. We filtered for primary alignments with alignment lengths > 10 kB and mapping quality > 40.

## RESULTS AND DISCUSSION

### Linkage map

Linkage map construction initially assigned 3,182 SNPs to 36 linkage groups in the pre-framework map. After expanding linkage analysis to the full SNP dataset, we assigned 12,151 SNPs to 41 linkage groups. We proceeded to build linkage maps for the 34 linkage groups that contained more than 5 markers (Figure 2). Our framework map with marker order supported by LOD > 5 consists of 4,468 SNPs with a total sex-averaged autosomal genetic map length of 2446.78 cM and mean genetic distance between markers of 0.56 cM (± 1.33 cM). The female and male autosomal map lengths were 2373.56 cM and 2567.09 cM, respectively (Figure 2). We include the full linkage map in Table S2.

**Figure 2.**
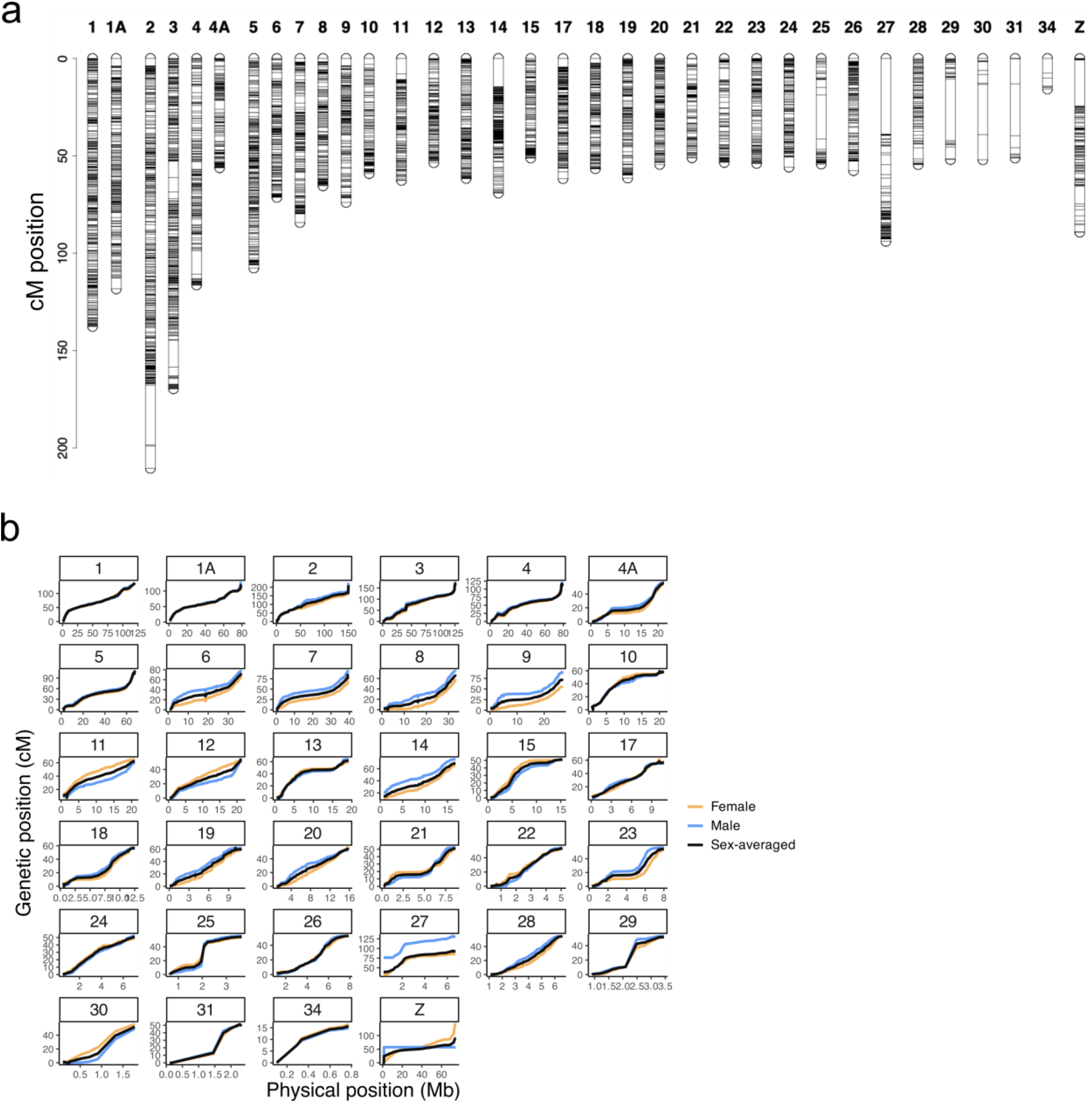
A genetic linkage map for the Florida Scrub-Jay. (A) Each column represents a linkage group (chromosome). Black tick marks represent marker locations of the LOD5 framework map, with linkage group lengths measured in Kosambi cM. This plot was created with the R package *LinkageMapView* (Ouellette *et al*. 2018). (B) Sex-averaged and sex-specific linkage maps for the Florida Scrub-Jay, with genetic position (Kosambi cM) shown relative to physical location (Mb). Please see Table S2 for the full linkage map.

### Genome sequencing and assembly

PacBio HiFi long-read sequencing yielded 84 Gb of raw read data, with a mean read length of 14.55 Kb. We created 4 draft assemblies with hifiasm: primary, alternate, maternally-resolved haplotype, and paternally-resolved haplotype. We moved forward with the primary assembly, as it was the most contiguous (L50/N50 of 18 contigs/17.7 Mb) and had the highest BUSCO scores of the three draft assemblies (97.1% completeness; Table S1). Scaffolding with SALSA and Juicer generated an assembly with 699 scaffolds and an L50/N50 of 9 scaffolds/33.36 Mb (Figure S1). Linkage map-aided scaffolding with ALLMAPS identified 34 linkage groups. Of these linkage groups, 31 (including the Z) were associated with complete chromosomes previously identified in v2 of the Florida Scrub-Jay genome (Driscoll & Beaudry, *et al*. 2021). The remaining 3 linkage groups were newly-assembled chromosomes with sequence homology to chromosomes 30, 31, and 34 in the Zebra finch (Figure 3). Our whole genome assembly displayed broad mapping synteny with other bird genomes, with 81%, 81.5%, 83.7%, and 42.2% of the Florida Scrub-Jay genome aligned to Zebra finch, New Caledonian crow, California Scrub-Jay, and chicken, respectively. As expected, the whole-genome alignment with the most distantly related chicken yielded the most rearrangements and sequence differences, while the alignment with the congeneric California Scrub-Jay yielded the fewest (Figure S2).

**Figure 3.**
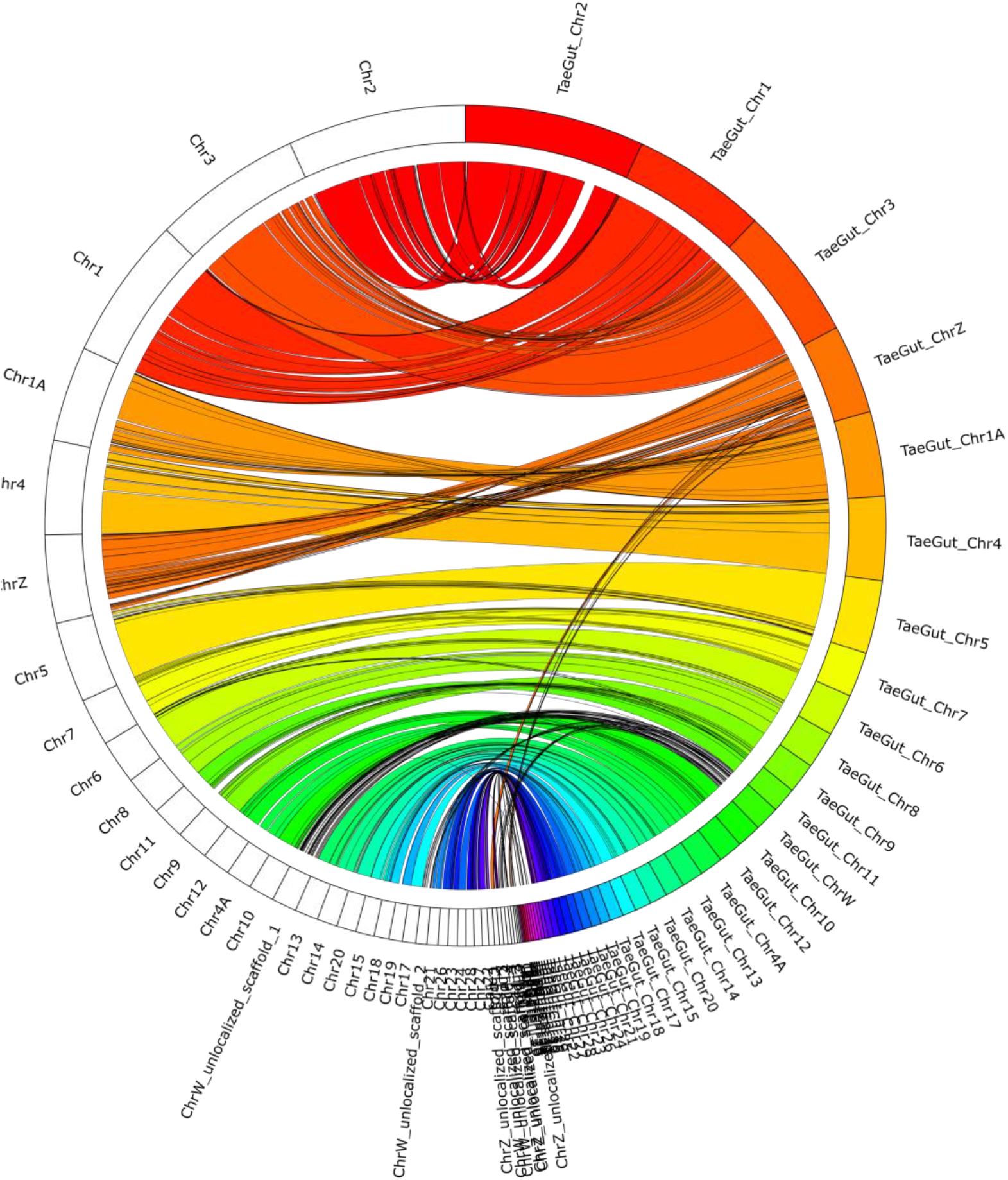
Sequence homology between the Florida Scrub-Jay and the Zebra finch (*Taeniopygia guttata*; bTaeGut1.4.pri). The outer ring represents genome sequence divided into chromosomes/scaffolds: white bars on the left represent Florida Scrub-Jay scaffolds and colored bars on the right represent Zebra finch scaffolds, with colored ribbons showing sequence alignment. For clarity, we filtered for alignments > 100kb. We created these plots with Circos v. 0.69-9 (Krzywinski *et al*. 2009) with code adapted from the online tutorial https://bioinf.cc/misc/2020/08/08/circos-ribbons.html.

Next, we identified sex-linked scaffolds by comparing average read coverage per scaffold across 25 male and 25 female Florida Scrub-Jays. We found 17 scaffolds that significantly differed in coverage between males and females. Due to their small size, we were only able to confirm 7 of these scaffolds, 3 Z-linked and 4 W-linked, as homologous to Zebra finch sex chromosome sequence (Figure 3, 4). The largest Z-linked scaffold, which corresponds to the Z linkage group, aligned to ∼99% of the Zebra finch Z chromosome (Figure 3). The largest W-linked scaffold was equal in size to the entire Zebra finch W chromosome (∼21 Mb), but the 3 additional W-linked scaffolds added 15.8 Mb in sequence, yielding a total of 36.8 Mb of sequence identified as part of the Florida Scrub-Jay W chromosome (Figure 3). Finally, we mapped all W-linked scaffolds onto the paternally-resolved haplotype assembly. After filtering using the scheme described previously, we yielded no alignments, further confirming the W assignment of these scaffolds. At the end of these analyses, we labeled the largest Z-linked scaffold as the Z chromosome, the 2 additional Z-linked scaffolds as unlocalized Z sequence, and the 4 W-linked scaffolds as unlocalized W sequence.

**Figure 4.**
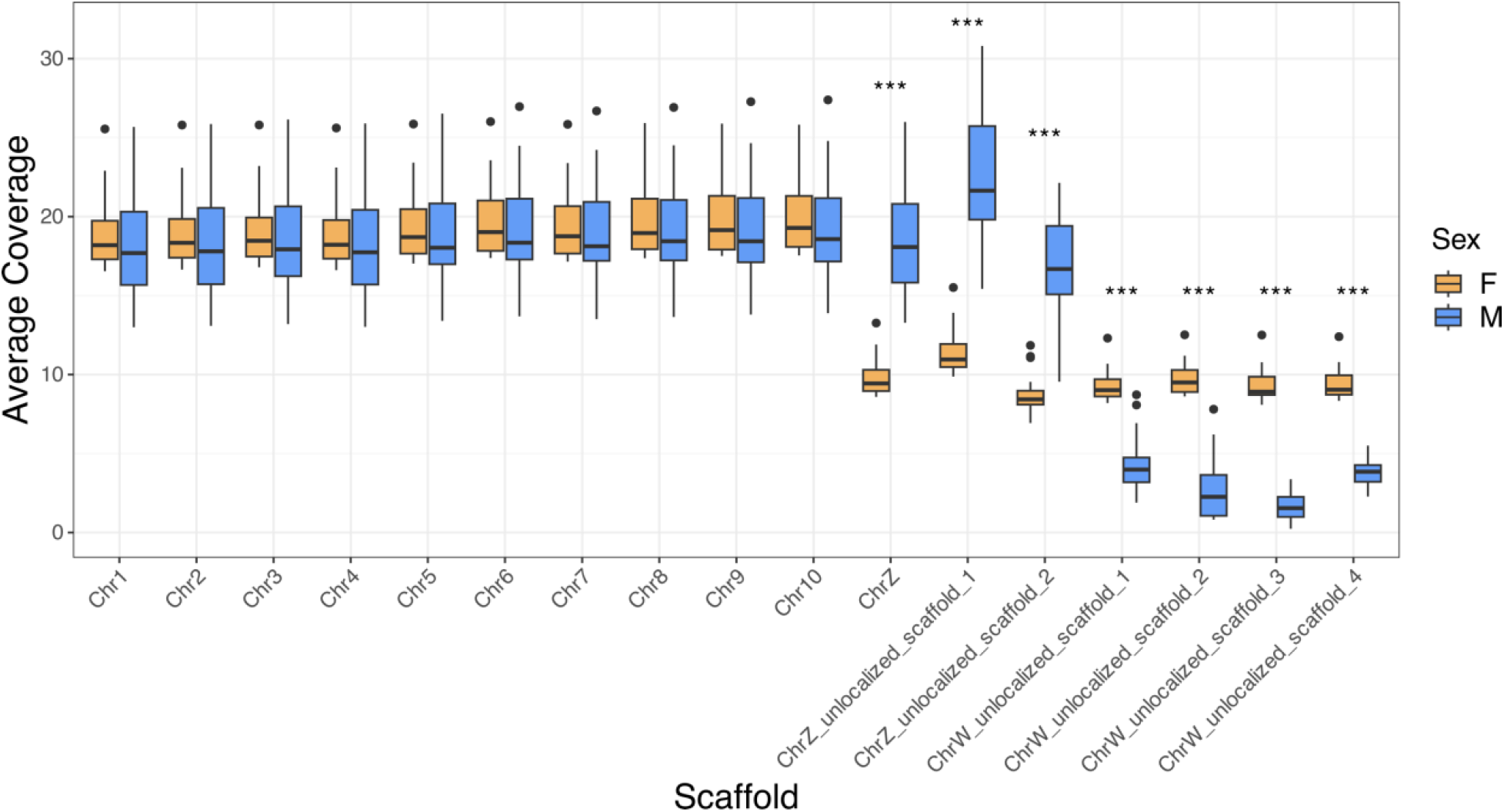
Average read depth of sex-linked scaffolds in 25 female (yellow) and male (blue) Florida Scrub-Jays. We include the first 10 autosomes for comparison. W-linked scaffolds show approximately twice the coverage in females when compared to males, while Z-linked scaffolds show approximately twice the coverage in males when compared to females. Scaffolds with significantly different average read depths from a t-test are indicated as follows: * = p < 2.26 × 10^-5^, ** = p < 2.26 × 10^-6^, *** = p < 2.26 × 10^-7^ (p-values were Bonferroni-corrected for 442 comparisons).

During decontamination screening with Blobtoolkit, one scaffold was identified as belonging to a non-Chordate (specifically, to *Drosophila melanogaster* in Arthropoda; Figure S3); however, we believe this result is a computational artifact because the BLAST hits had low percent identity (*pident* = 76-77%) and represented a very low percentage (∼3.3%) of the total scaffold length. We therefore retained this scaffold in the final genome assembly. Our final genome assembly is 1.32 Gb long and consists of 660 scaffolds, 87.6% of which belonged to 33 named autosomes and the sex chromosomes, with an L50/N50 of 7 scaffolds/88.05 Mb, an average depth of 63x, and a BUSCO completeness score of 97.1% with respect to the aves_odb10 database (Table 1). These quality measures are similar to those of other published genome assemblies in Corvidae and Passeriformes (Table 2). The Florida Scrub-Jay has the longest genome amongst the species considered, and is comparable in length to the more closely-related California Scrub-Jay (1.35 Gb; DeRaad *et al*. 2023).

**Table 1.**
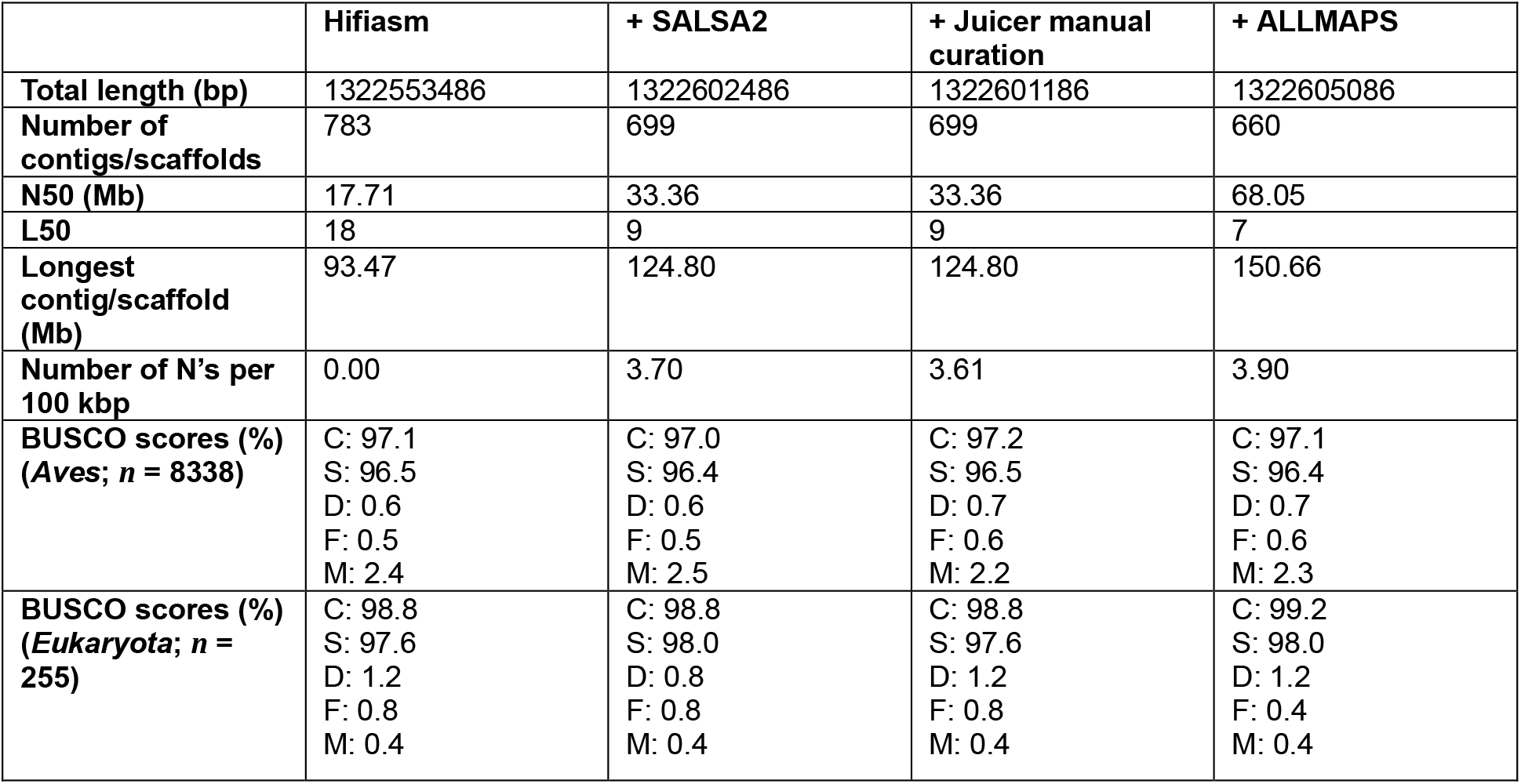
Basic assembly statistics for each step of the v3 Florida Scrub-Jay genome assembly. The Hifiasm column reports statistics for contigs, while all other columns report statistics for scaffolds. The *Aves* row of BUSCO (Benchmarking Universal Single-Copy Orthologs) scores uses the aves_odb10 (2024-01-08) database with 8338 BUSCOs available. The *Eukaryota* row of BUSCO scores uses the eukaryote_odb10 (2024-01-08) database with 255 BUSCOs available. BUSCO parameters are as follows: C: Complete, S: Complete and single-copy, D: Complete and duplicated, F: Fragmented, M: Missing (Manni *et al*. 2021).

**Table 2.** Summary statistics for the Florida Scrub-Jay (*A. coerulescens*) genome and annotation compared to other similar bird species in Corvidae and Passeriformes (New Caledonian crow, *Corvus moneduloides*; Hawaiian crow, *Corvus hawaiiensis*; Hooded crow, *Corvus cornix*; Collared fly catcher, *Ficedula albicollis*; Zebra finch, *Taeniopygia gutatta*). We calculated annotation statistics for each species by inputting their publicly available annotation files (GFF) into the AGAT toolkit script *agat_sp_statistics.pl* (Dainat 2023). Note that genome assembly length, contig N50, and genome BUSCO scores are genome summary statistics, while all other statistics are gene annotation summary statistics. BUSCO parameters are as follows: C: Complete, S: Complete and single-copy, D: Complete and duplicated, F: Fragmented, M: Missing (Manni *et al*. 2021). Modified from (Peona *et al*. 2023).

|  | Florida Scrub-Jay | New Caledonian crow | Hawaiian crow | Hooded crow | Collared flycatcher | Zebra finch |
| --- | --- | --- | --- | --- | --- | --- |
| Genome assembly length (Gb) | 1.3 | 1.1 | 1.2 | 1.0 | 1.1 | 1.22 |
| Contig N50 (Mb) | 17.7 | 11.5 | 23.1 | 8.6 | 0.41 | 0.038 |
| No. of genes | 17,812 | 16,167 | 16,414 | 14,435 | 16,763 | 17,561 |
| Mean gene length (bp) | 23,127 | 36,436 | 35,242 | 37,961 | 31,394 | 26,458 |
| Number of CDS | 26,521 | 43,047 | 43,147 | 36,899 | 16,763 | 17,561 |
| Mean length of CDS (bp) | 1,912 | 2,293 | 2,332 | 2,275 | 1,942 | 1,677 |
| No. of exons | 311,707 | 638,785 | 648,529 | 555,077 | 189,043 | 171,767 |
| Mean exon length (bp) | 162 | 291 | 283 | 293 | 253 | 255 |
| Mean no. exons per gene | 11.8 | 14.1 | 14.3 | 14.3 | 12.2 | 10.3 |
| No. of introns | 285,186 | 563,402 | 573,808 | 490,050 | 171,236 | 153,909 |
| Genome BUSCO scores (%) (Aves; *n* = 8338) | C: 97.1<br>S: 96.4<br>D: 0.7<br>F: 0.6<br>M: 2.3 | C: 96.8<br>S: 96.3<br>D: 0.5<br>F: 0.5<br>M: 2.7 | C: 97.3<br>S: 96.8<br>D: 0.5<br>F: 0.5<br>M: 2.2 | C: 94.8<br>S: 94.4<br>D: 0.4<br>F: 0.6<br>M: 4.6 | C: 96.5<br>S: 96<br>D: 0.5<br>F: 0.8<br>M: 2.7 | C: 93.8<br>S: 91.9<br>D: 1.9<br>F: 2.3<br>M: 3.9 |

### Genome annotation

*Repetitive content.* We identified 247 Mb of interspersed repeats throughout the genome, comprising 18.71% of the total genome length (Figure 5A). Repetitive content of the v3 genome was more than double that of the v1 and v2 assemblies, which both had an estimated interspersed repeat content of 8.7% (Table S4). The new v3 assembly added ∼155 Mb of identified repetitive content and yielded more long interspersed nuclear elements (LINEs), long tandem repeats (LTRs), non-LTR retroelements (*e.g.*, short interspersed nuclear elements (SINEs)), satellite sequences, simple repeats, and unclassified repetitive elements (Figure S4A). This increase in repetitive content between our long-read (v3) and short-read (v2) genome assemblies matches similar patterns observed in sparrows (Benham *et al*. 2023b). Avian genomes have long been thought to have low repeat content (< 10%) (Ellegren 2010; Zhang *et al*. 2014b), but genomes assembled with new long-read sequencing technologies are indicating that repeat content in bird genomes has previously been underestimated.

**Figure 5.**
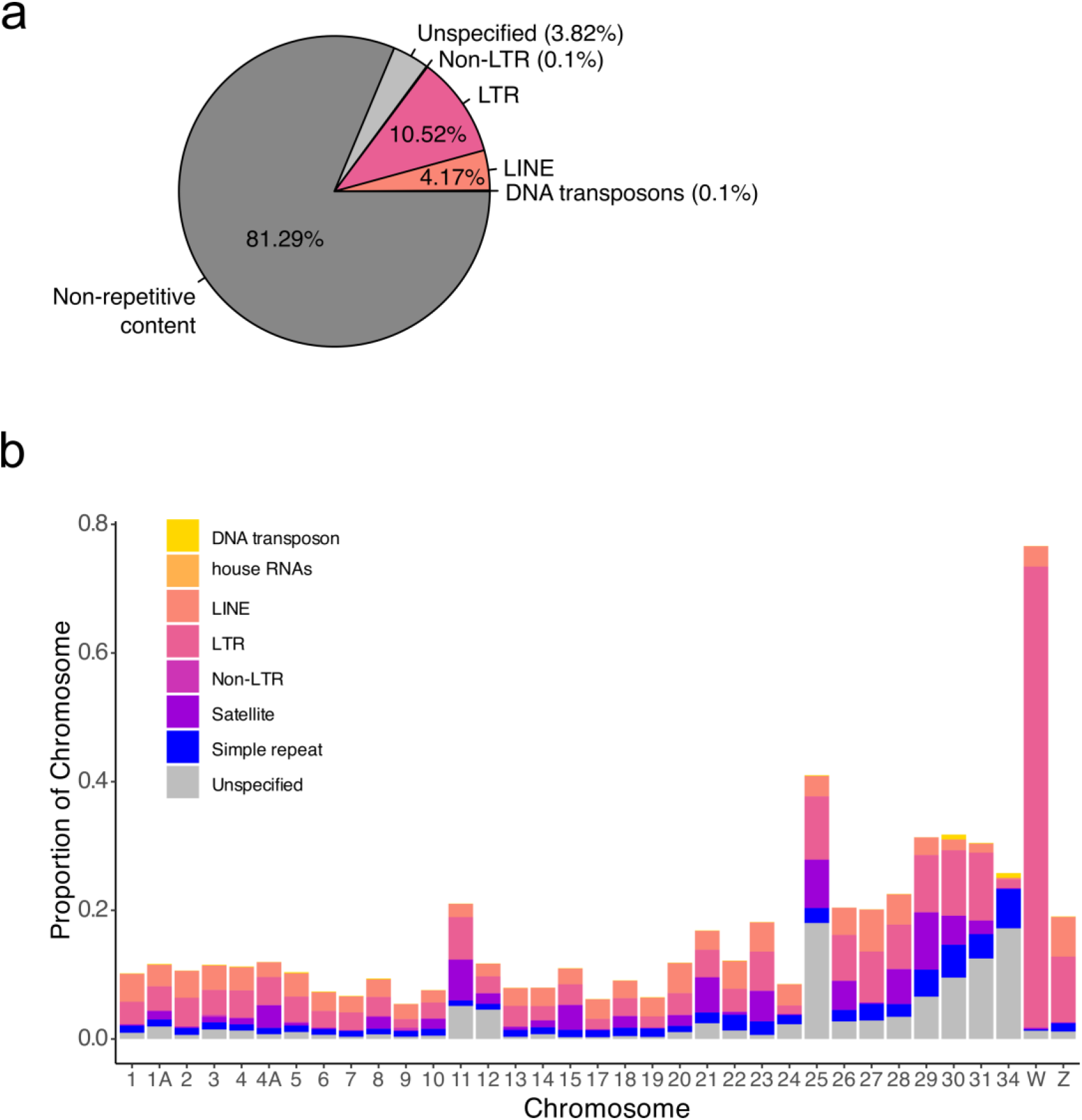
Repeat annotation of the Florida Scrub-Jay genome. (A) Summary of interspersed repeat content across the whole genome. (B) Repetitive content of each chromosome, colored by transposable element superfamily (Kapitonov and Jurka 2008).

The median length of all TE superfamilies differed significantly between the v2 and v3 assemblies: long-read sequencing assembled, on average, longer house RNA, satellite, and simple repeat elements (Figure S4B). These results highlight the effectiveness of long-read sequencing technology in assembling both shorter and longer repetitive elements. The most common element across the genome was LTR retrotransposons (14.79%) followed by LINEs (4.17%) (Figure 5), aligning with patterns seen in other avian genomes, particularly amongst songbirds (Kapusta and Suh 2017; Boman *et al*. 2019; Weissensteiner *et al*. 2020). The W-linked scaffold was a strong outlier in repetitive content, with 76.6% of the sequence characterized as interspersed repeats spanning the entire chromosome (Figure S5). The majority of these elements were classified as LTRs, supporting the role of the W chromosome as a haven for repetitive content, particularly long LTR elements, in birds (Peona *et al*. 2021a, 2021b).

*Gene content.* We annotated a total number of 17,812 genes throughout the genome, with a mean gene length of 23.1 Kb and a mean of 11.8 exons per gene (Table 2). Of the identified genes, 92.5% were annotated with functional information and 84.3% had an associated gene ontology term. BUSCO completeness of the transcriptome was 95.5% for the *Aves* database and 98.4% for the *Eukaryota* database. Mean gene length, mean number exons per gene, and all other annotation quality measures are comparable to that of other similar avian species (Table 2).

Notably, we annotated the greatest number of genes (17,812) amongst the species considered (Table 2).

## CONCLUSION

We report a high-quality genome assembly, associated annotation, and a linkage map for the Florida Scrub-Jay. Using a combination of long-read sequencing, Hi-C data, and our linkage map, we generated a highly contiguous genome assembly, with a size of 1.32 Gb, an N50 of 88.05 Mb, and a BUSCO completeness score of 97.1% (Table 1). Additionally, we provide the first assembly of the W chromosome in this species, as well as three newly identified chromosomes. This annotated genome assembly and linkage map will facilitate more detailed genetic analyses, such as the exploration of haplotype dynamics across space and time and the genetic architecture of fitness, and open the door for new and exciting questions about the biology, ecology, and evolution of this Federally Threatened species.

## Supporting information

Linkage map

## DATA AVAILABILITY

The genome assembly, annotation, and associated raw data for this project are available on NCBI: accession X (pending), BioProject PRJNA1076903, BioSample SAMN39956395. All associated code and data are available at github.com/faye-romero/FSJ-genome.

## ACKNOWLEDGMENTS

Reed Bowman (1958-2023; Avian Ecology Program, Archbold Biological Station, Venus, FL 33960) was a co-author on this project. We would like to thank the Center for Integrated Research Computing at the University of Rochester for their technical expertise, Tim Sackton for sharing advice and comparative genomic resources, Emiliano Martí for assistance with the genome annotation, and the Chen lab for additional support.

## FUNDING

Funding provided by the National Science Foundation (NSF) (grant DEB-1257628) and the Cumming Foundation. F.G.R. was supported by a National Institute of Health (NIH) grant to N.C. (1R35GM133412) and a NSF Graduate Research Fellowship (DGE-1939268). F.E.G.B. was supported by the aforementioned NIH grant to N.C. and a NSF Postdoctoral Research Fellowship (2109639).

## CONFLICT OF INTEREST

The authors declare no competing interests.

## Notes

### Competing Interest Statement

The authors have declared no competing interest.

