## Supplemental Material for "A new high-quality genome assembly and annotation for the threatened Florida Scrub-Jay (*Aphelocoma coerulescens*)"

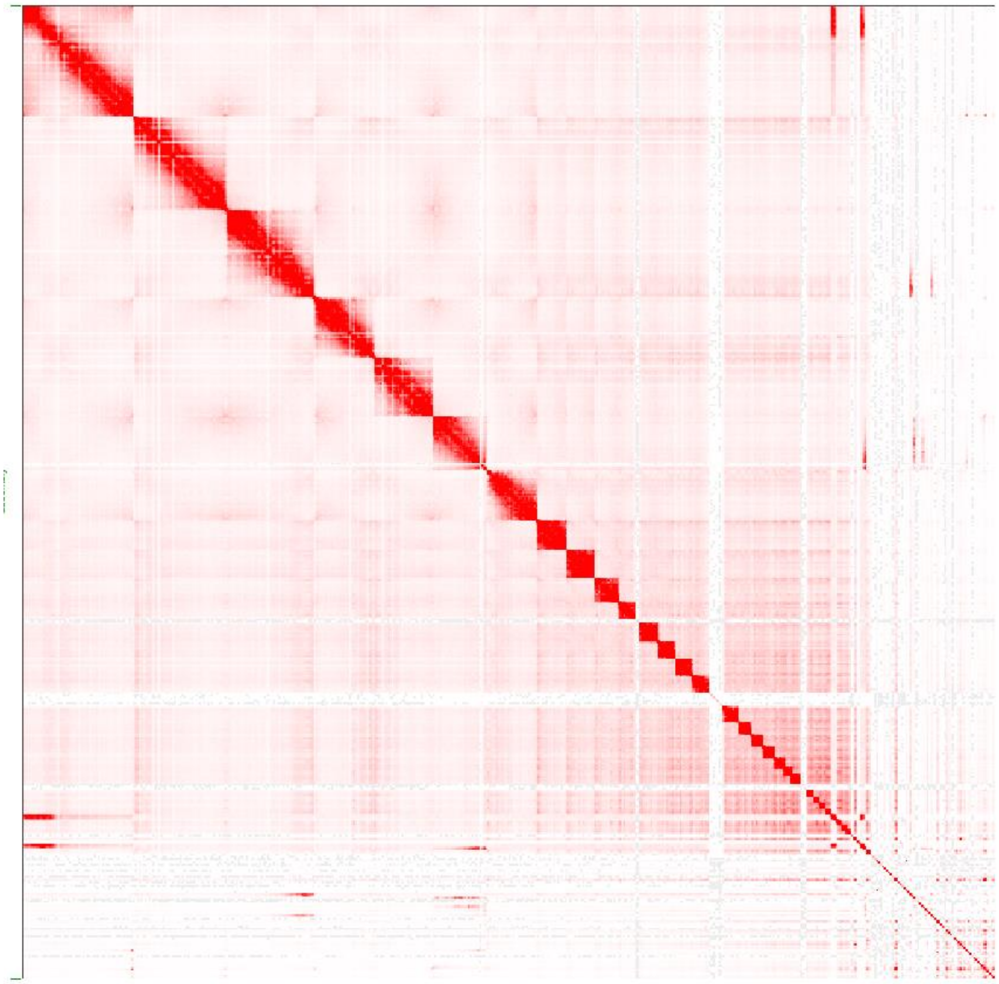

**Figure S1.** Hi-C contact map of the final genome assembly. We created this map using the Juicer/JuiceBox suite (Durand *et al.* 2016).

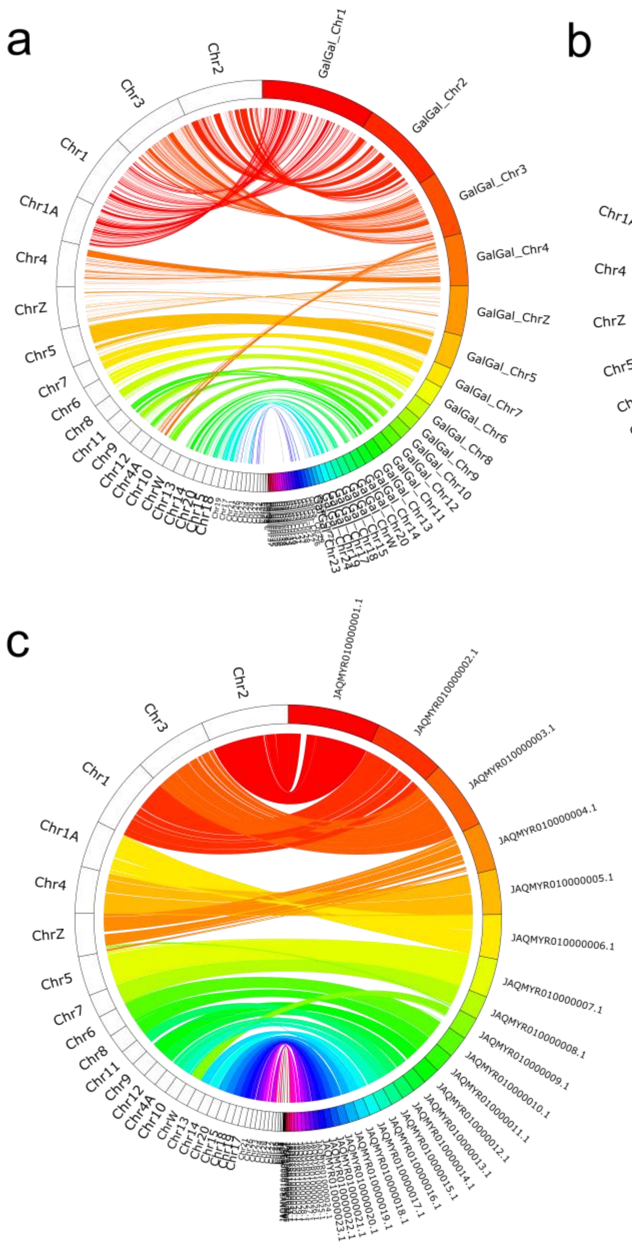


**Figure S2.** Sequence homology between the v3 Florida Scrub-Jay genome assembly and the (a) chicken, (b) New Caledonian crow, and (c) California Scrub-Jay genomes. The outer ring represents genome sequence divided into chromosomes/scaffolds: white bars on the left represent Florida Scrub-Jay scaffolds and colored bars on the right represent the reference species sequence, with colored ribbons showing sequence alignment. The California Scrub-Jay reference genome did not have named chromosomes; we therefore sorted the whole genome by descending scaffold size and extracted the 35 largest scaffolds for alignment. For Florida Scrub-Jay chromosomes with multiple unlocalized scaffolds, only the largest scaffold was used for alignment. For clarity, we filtered the chicken and New Caledonian crow plots for alignments > 100kb and the California Scrub-Jay plot for alignments > 800kb. We created these plots with Circos v. 0.69-9 (Krzywinski *et al.* 2009) with code adapted from the online tutorial https://bioinf.cc/misc/2020/08/08/circos-ribbons.html.

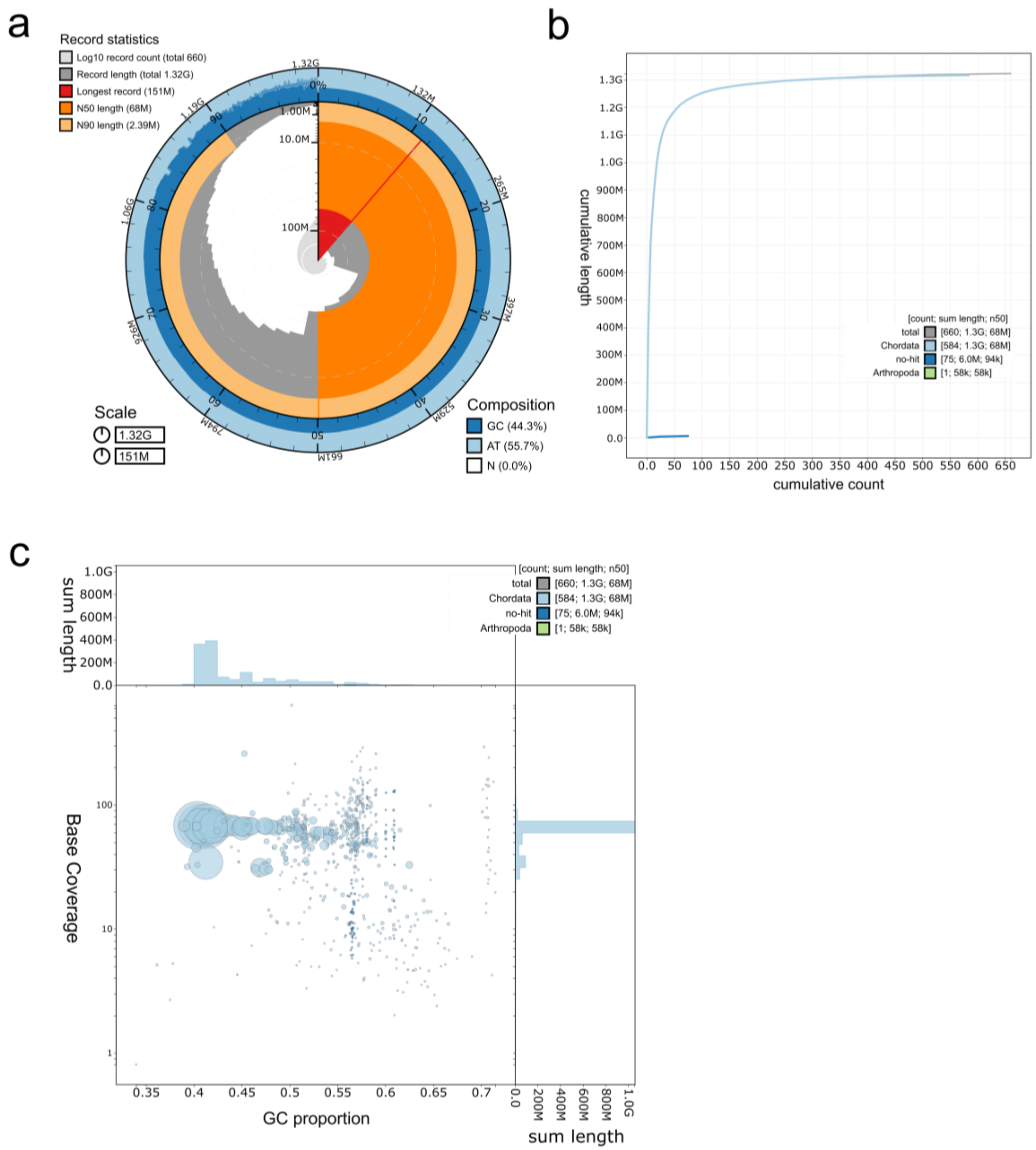


**Figure S3.** Decontamination screening results. (a) Snail plot describing summary statistics of the v3 genome assembly. (b) Plot displaying cumulative length of sequences assigned to different taxonomic categories. (c) Blob plot of base coverage across the genome (y axis) against proportion of GC content across the genome (x axis). Histograms show the distribution of sequence lengths along each axis. We viewed and created all plots with the BlobToolKit pipeline (Challis *et al.* 2020).

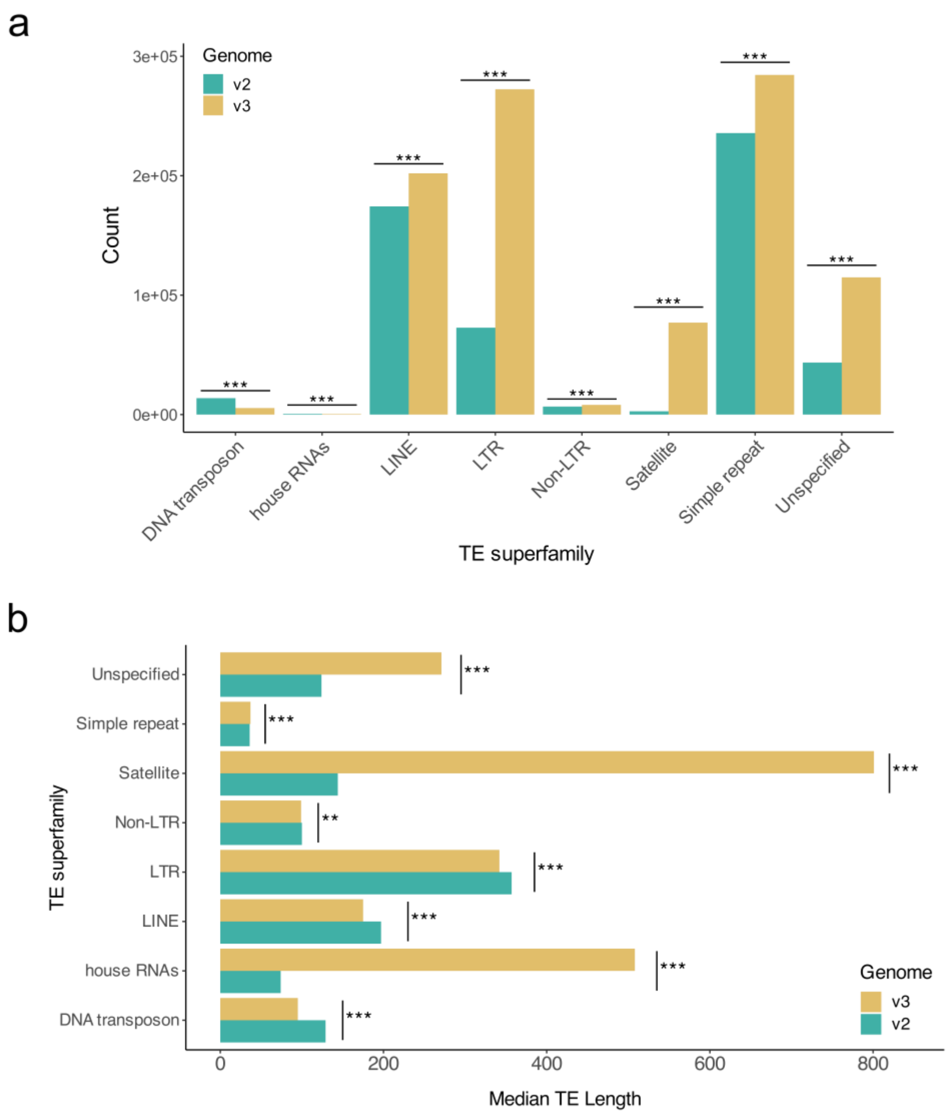


**Figure S4.** Comparison of repetitive element annotation between Illumina short-read (v2; Driscoll and Beaudry *et al.* 2021) and PacBio HiFi long-read (v3; presented here) Florida Scrub-Jay genome assemblies. (a) Number of repetitive elements annotated across each genome, separated by transposable element (TE) superfamily (Kapitonov and Jurka 2008). TE counts for each superfamily differed significantly between v2 and v3 (Proportion test; $p < 0.001$ for all). (b) Median length of TEs annotated across each genome, separated by TE superfamily (Kapitonov and Jurka 2008). Median TE length for each superfamily differed significantly between v2 and v3 (Wilcoxon rank sum test; $p < 0.01$ for all). Significance codes are as follows: * = $p < 0.05$, ** = $p < 0.01$, *** = $p < 0.001$.

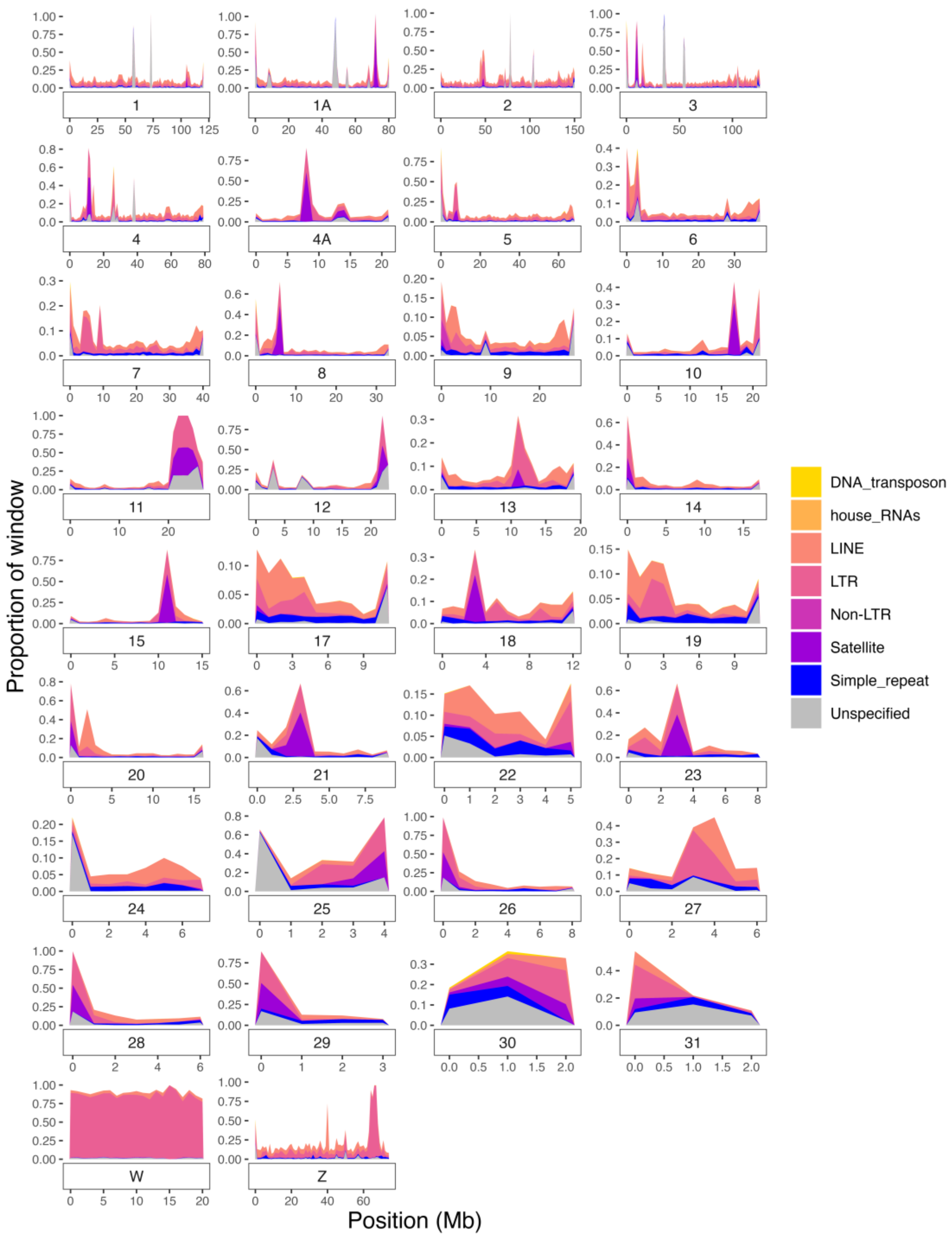


**Figure S5.** Transposable element composition across 1 Mb windows for each chromosome. Note that the y-axis range varies across chromosomes. Average repeat content for the W chromosome (76.6%) is higher than that for the other chromosomes (16%). We grouped transposable elements by superfamily (Kapitonov and Jurka 2008).

| Purpose | Software | Version | Options |
|---|---|---|---|
| *De novo genome assembly* | | | |
| Remove adapter sequences from PacBio HiFi raw reads | Cutadapt | 2.3 | --discard-trimmed --overlap=35 -b ATCTCTCTCAACAACAACAACGGA GGAGGAGGAAAAGAGAGAGAT -b ATCTCTCTCTTTTCCTCCTCCTCCG TTGTTGTTGTTGAGAGAGAT |
| Genome assembly | Hifiasm (HiFi-only assembly mode) | 0.16.1 | default |
| Trim and filter parental short reads | fastp | 0.21.0 | default |
| Merge parental paired-end short reads | PEAR | 0.9.11 | default |
| Build k-mer hash tables for parental short reads | yak | 0.1 (r56) Commit: 76db8e8 | count -k31 -b37 -t16<br>count -k31 -b37 -t16 |
| Genome assembly | Hifiasm (Trio-binning assembly mode) | 0.16.1 | default |
| Map Hi-C reads to draft genome assembly | Arima Genomics Mapping Pipeline | February 8, 2019 release (Document Part Number A160156 v01) | default |
| Scaffold assembly | SALSA | 2.3 | -e GATC -i 5 -m yes |
| Scaffold and visualize assembly | Juicer | 1.6 | *juicer.sh*<br>-s DpnII -C 180000000 --assembly<br><br>*run-assembly-visualizer.sh*<br>default<br><br>*run-asm-pipeline-post-review.sh*<br>--sort-output -r |
| Scaffold assembly | ALLMAPS (JCVI Utilities Libraries) | 1.3.7 | -m jcvi.assembly.allmaps merge<br>-m jcvi.assembly.allmaps path |
| Decontamination screening | BlobToolKit | 3.1.0 | add --busco<br>add --hits --taxrule bestsumorder<br>add --cov |
| Decontamination screening | BLAST | 2.10.0+ | -outfmt 6 |
| Whole genome alignments | Minimap2 | 2.26 | -x asm20 |
| Plot whole genome alignments | Circos | 0.69-9 | default |
| *Sex chromosome identification* | | | |
| Trim and filter Illumina raw reads | Trim Galore | 0.6.10 | --paired --fastqc |
| Map raw reads to genome assembly | BWA-MEM | 0.7.4 | default |
| Filter reads for quality | Samtools | 1.9 | view -b -f 0x2 -F 260 -q 20 |

| | | | |
|---|---|---|---|
| Mark and remove duplicate reads | Sambamba | 1.0.1 | markdups --remove-duplicates |
| Filter reads for quality | Bamtools | | filter -tag NM:<=2 |
| Calculate read depth | Samtools | 1.9 | mpileup --positions --fasta-ref |
| *Genome annotation* | | | |
| Annotate *de novo* repeat families | RepeatModeler | 2.0.4 | -LTRStruct |
| Identify and classify repetitive elements | RepeatMasker | 4.1.4 | -gff -s -a -xsmall |
| Align RNA-seq reads to genome assembly | STAR | 2.7.3 | --twopassMode Basic --readFilesCommand zcat --outSAMstrandField intronMotif |
| Assign read groups to RNA-seq reads | Picard | 2.27.4 | AddOrReplaceReadGroups -VALIDATION_STRINGENCY LENIENT -RGPL RNA |
| Gene annotation | BRAKER3 | 3.0.6 | --prot_seq --rnaseq_sets_ids --gff3 |
| Assign functional information to gene annotation | InterProScan | 5.66-98.0 | --goterms --iprlookup |
| Gene annotation | BLAST | 2.10.0+ | blastp -evalue 0.000001 -outfmt 6 |
| Merge gene annotations, calculate summary statistics | AGAT | 1.2.0 | *agat_sp_manage_functional_annotation.pl*<br>--gff3 --blast --db --interpro --pcds<br><br>*agat_sp_statistics.pl*<br>--gff --gs 1322604086 -d<br><br>*agat_sp_functional_statistics.pl*<br>--gff --gs 1322604086 |

**Table S1.** Descriptions of programs and software used during genome assembly and annotation.

Any options listed are in addition to default parameters.

**Table S2**: A framework (LOD > 5) linkage map for the Florida Scrub-Jay constructed using CRI-MAP v. 2.507 (Green *et al.* 1990). We present the physical locations of each marker in the v3 genome assembly and genetic positions for the sex-averaged, female, and male linkage maps in Kosambi cM.

[File: *FileS1.tsv*]

| | Primary (HiFi-only mode) | Alternate (HiFi-only mode) | Maternally-resolved haplotype (Trio-binning mode) | Paternally-resolved haplotype (Trio-binning mode) |
|---|---|---|---|---|
| **Total length (bp)** | 1322553486 | 984436143 | 1293735984 | 738092893 |
| **Number of contigs/scaffolds** | 783 | 4851 | 990 | 1133 |
| **N50 (Mb)** | 17.71 | 0.95 | 13.39 | 5.35 |
| **L50** | 18 | 280 | 25 | 34 |
| **Longest contig/scaffold (Mb)** | 93.47 | 6.53 | 81.63 | 34.17 |
| **Number of N's per 100 kbp** | 0.00 | 0.00 | 0.00 | 0.00 |
| **BUSCO scores (*Aves* database)** | C: 97.1%<br>S: 96.5%<br>D: 0.6%<br>F: 0.5%<br>M: 2.4% | C: 76.1%<br>S: 75.1%<br>D: 1.0%<br>F: 0.9%<br>M: 23.0% | C: 92.8%<br>S: 87.7%<br>D: 5.1%<br>F: 0.5%<br>M: 6.7% | C: 49.4%<br>S: 48.3%<br>D: 1.1%<br>F: 0.5%<br>M: 50.1% |

**Table S3.** Assembly statistics for each draft assembly created with hifiasm v. 0.16.1. BUSCO parameters are as follows: C: Complete, S: Complete and single-copy, D: Complete and duplicated, F: Fragmented, M: Missing (Manni *et al.* 2021).

| | Version 1 | Version 2 | Version 3 |
|---|---|---|---|
| *Assembly statistics* | | | |
| **Total length (bp)** | 1060889110 | 1060969718 | 1322605086 |
| **Number of contigs/scaffolds** | 1678 | 878 | 660 |
| **N50 (Mb)** | 7.66 | 74.35 | 68.05 |
| **L50** | 40 | 5 | 7 |
| **Longest contig/scaffold (Mb)** | 29.17 | 156.03 | 150.66 |
| **Number of N's per 100 kbp** | 1907.36 | 1914.34 | 3.90 |
| **BUSCO scores (%) (*Aves* database)** | C: 97.1<br>S: 96.7<br>D: 0.4<br>F: 0.5<br>M: 2.4 | C: 97.2<br>S: 96.8<br>D: 0.4<br>F: 0.5<br>M: 2.3 | C: 97.1<br>S: 96.4<br>D: 0.7<br>F: 0.6<br>M: 2.3 |
| *Repetitive element annotation statistics (% of genome)* | | | |
| **Total interspersed repeats** | 8.72 | 8.76 | 18.71 |
| **Retroelements** | 7.69 | 7.74 | 14.79 |
| **LTR elements** | 3.16 | 3.21 | 10.52 |
| **DNA transposons** | 0.34 | 0.34 | 0.1 |
| **Satellites** | 0.09 | 0.07 | 3.37 |
| **Unclassified** | 0.7 | 0.68 | 3.82 |

**Table S4.** Assembly and repetitive element annotation statistics for 3 versions of the Florida Scrub-Jay genome: Version 1 (Feng *et al.* 2020), Version 2 (Driscoll and Beaudry *et al.* 2021), and Version 3 (presented here). BUSCO parameters are as follows: C: Complete, S: Complete and single-copy, D: Complete and duplicated, F: Fragmented, M: Missing (Manni *et al.* 2021).
